# A panel of *CYP6Z* genes drives broad-spectrum cross-resistance to public health insecticides in the major malaria vector *An. gambiae* s.s.

**DOI:** 10.64898/2025.12.22.695880

**Authors:** Mersimine F. M. Kouamo, Arnaud Tepa, Abdullahi Muhammad, Vanessa B. Ngannang-Fezeu, Jonas A. Kengne-Ouafo, Murielle Wondji, Helen Irving, Sulaiman S. Ibrahim, Charles S. Wondji

## Abstract

The spread of multiple and intense insecticide resistance in major African malaria vectors is jeopardising control efforts. Mitigating this threat requires deciphering its underlying molecular mechanisms. Here, by integrating transcriptomic profiling, analysis of genetic diversity and functional genomics approaches, we established the role of the *CYP6Z* genes in conferring multiple- and cross-resistance to insecticides.

Investigation of *CYP6Z* gene expression and genetic diversity analyses indicate that resistance is primarily associated with transcriptional upregulation rather than fixed coding mutations since no predominant haplotype was selected. Structural characterisation reveals a flexible, promiscuous active site in CYP6Z enzymes, enabling binding of multiple insecticide classes. *In vitro* functional validation confirms that recombinant *CYP6Z3* efficiently metabolizes deltamethrin (percentage depletion of 59%), permethrin (52%), α-cypermethrin (37%), pirimiphos-methyl (36%), fenitrothion (25%), propoxur (25%), and possibly bendiocarb (17%). RNAi-mediated knockdown of *CYP6Z* genes in field-collected *An. gambiae s.s.* restores susceptibility to several insecticides including clothianidin: dsCYP6Z1 (mortality = 50.29%; *p* < 0.01), dsCYP6Z2 (47.28%; *p* < 0.01) and dsCYP6Z3 (39.12%; *p* < 0.05) compared to the control (mortality = 27.67%). Furthermore, transgenic expression in *Drosophila melanogaster* flies revealed that expression of *CYP6Z* genes alone confer cross-resistance to pyrethroids, organophosphates, carbamates and neonicotinoids, but increases susceptibility to the pro-insecticide chlorfenapyr: *CYP6Z1* (mortality = 100%; p < 0.01), *CYP6Z3* (98.18%; *p* < 0.01), and *CYP6Z4* (98.01%; *p* < 0.01) vs. control (88.03%).

This study establishes *An. gambiae CYP6Z* genes operate in concert to drive a broad-spectrum metabolic resistance, towards several classes of public health insecticides, while in contrast bioactivating chlorfenapyr to its more insecticidally active metabolite.

## Introduction

The widespread use of insecticides in malaria vector control has led to the emergence and rapid spread of resistance in major malaria vectors such as *An. gambiae s.s.* and *An. coluzzii*, threatening the efficacy of control tools such as long-lasting insecticidal nets (LLINs) and indoor residual spraying (IRS) (Ranson and Lissenden, 2016; WHO, 2024). Among the various mechanisms of insecticide resistance, metabolic resistance represents a serious challenge to the success of malaria vector control (Hemingway et al., 2002). By enhancing the activity and/or expression of detoxification enzymes, malaria vectors are able to degrade or sequester insecticides before they can reach their intended neuronal targets (Hemingway et al., 2004; Ranson et al., 2002). This mechanism, unlike target-site resistance, could confer broad-spectrum protection across multiples insecticide classes and can evolve rapidly under insecticide pressure (Schuler and Berenbaum, 2013) Among the major classes of detoxification enzymes, including the P450 monooxygenases (P450s), carboxylesterases, and glutathione S-transferases (*GSTs*), the P450s are the most frequently implicated in high-level resistance to pyrethroids (Li et al., 2007; Tepa et al., 2022; Vontas et al., 2018). In *An. gambiae s.s.*, P450s including *CYP6P3*, *CYP6M2*, *CYP6Z1* and *CYP9K1* have been functionally validated for their ability to confer pyrethroids resistance (Chiu et al., 2008; Kengne-Ouafo et al., 2024; Muller et al.; Nauen et al., 2022; Stevenson et al., 2011).

The *Anopheles CYP6Z* gene family is particularly noteworthy due to its repeated association with insecticide resistance in malaria mosquitoes across Africa (Ibrahim et al., 2016; Irving et al., 2012; Tepa et al., 2025). For instance, transcriptional profiling showed that *CYP6Z1* is overexpressed in pyrethroid and DDT-resistant field populations of *Anopheles gambiae* from Côte d’Ivoire from (Müller et al., 2007) Kenya (David et al., 2005), and Cameroon (Kala-Chouakeu et al., 2022; Kengne-Ouafo et al., 2024; Tepa et al., 2022). Similarly, in *An. gambiae s.s.* and *An. coluzzii* from Senegal, *CYP6Z1* and *CYP6Z2* were transcriptionally upregulated in highly pyrethroid-resistant mosquitoes alongside the presence of voltage-gated sodium channel knockdown resistance (*kdr*) mutations (Gueye et al., 2020). Beyond pyrethroids, *CYP6Z1* has been functionally validated in Malawi *An. funestus* and shown to metabolize both pyrethroids and carbamates (Ibrahim et al., 2016). In addition, the expression of *CYP6Z1* has also been associated with reduced efficacy of long-lasting insecticidal nets (LLINs) in *An. coluzzii*, where sublethal pyrethroid exposure enhanced metabolic detoxification, compromising bed net performance (Main et al., 2018; Meiwald et al., 2022).

Functional validation experiments have elucidated distinct metabolic capabilities within this gene family. In *An. funestus* from Malawi, recombinant enzyme assays demonstrated that *CYP6Z1* metabolizes both pyrethroids and carbamates (Ibrahim et al., 2016), while *An. gambiae CYP6Z1* was shown to metabolizes DDT (Chiu et al., 2008), providing direct evidence of its role in cross-resistance between these insecticide classes. In contrast, *CYP6Z2*, although consistently overexpressed in resistant *An. gambiae* populations across Senegal, Burkina Faso, Côte d’Ivoire, Ghana, Nigeria, and Cameroon (Ibrahim et al., 2023; Ingham et al., 2014; Mohammed et al., 2021), seemed unable to metabolize pyrethroid directly, (McLaughlin et al., 2008). However, its distant paralog, *Aedes aegypti CYP6Z8* was shown to metabolize the primary products of pyrethroid metabolism, 3-phenoxybenzoic alcohol and 3-phenoxybenzaldehyde generated by carboxylesterases (Chandor-Proust et al., 2013). Other studies have shown that *An. gambiae CYP6Z2* can metabolize other insecticides, e.g., pyriproxyfen (Yunta et al., 2016) and its ortholog *An. coluzzii CYP6Z2* was established to confer marginal resistance towards 3-phenoxybenzoic alcohol and 3-phenoxybenzaldehyde (Ibrahim et al., 2023). This functional profile highlights *CYP6Z1* as a primary pyrethroid metabolizer, while *CYP6Z2* performs downstream detoxification steps.

Similarly, *CYP6Z3* has been found transcriptionally upregulated in resistant populations in Côte d’Ivoire, Benin, and Cameroon (Antonio-Nkondjio et al., 2016; Fadel et al., 2024; Ibrahim et al., 2023; Mitchell et al., 2014; Saizonou et al., 2024). In addition, a recent study showed that expression of *CYP6Z3* and cuticular proteins in *An. gambiae* are associated with cross-resistance to pyrethroids and organophosphates (Saizonou et al., 2024).

Recent studies have suggested potential cross-resistance mediated by P450s to other insecticide classes such as carbamates, organophosphates, and novel compounds including neonicotinoids and pyrroles (e.g., clothianidin and chlorfenapyr), which are being incorporated into next-generation vector control products (Oxborough et al., 2019). For example, the study by Ibrahim et al. (2024) showed that *Anopheles funestus* CYP9A and CYP9B recombinant proteins together can metabolize DDT (producing dicofol), bendiocarb, clothianidin, and chlorfenapyr (via bioactivation to tralopyril) (Ibrahim et al., 2024). Moreover, it has recently been shown that the duplicated cytochrome P450 enzymes *CYP6P9a* and *CYP6P9b* increase mosquito susceptibility to the pro-insecticide chlorfenapyr by metabolically converting it into its highly toxic active form, tralopyril, demonstrating how P450-mediated bioactivation can enhance the potency of chlorfenapyr and lead to increased insecticide susceptibility (Tchouakui et al., 2024). In addition, recent work in *An. coluzzii* from Benin, demonstrated that resistance to the neonicotinoids acetamiprid and imidacloprid is associated with significant upregulation of *CYP6Z1* and *CYP6Z2*, alongside *CYP6M2* and *CYP6P4* (Tchigossou et al., 2024).

Besides progress in understanding the role of individual *An. gambiae s.s. CYP6Z* genes in insecticide resistance, the combined contribution of this gene sub-family in mediating cross-resistance to both established and new insecticide classes remains poorly characterized. This study filled this critical knowledge gap by integrating transcriptomic profiling, gene expression analysis, population genetic diversity assessment, *in silico* modelling/docking simulations, *in vitro* recombinant protein metabolism assays, and *in vivo* functional validation using transgenic *Drosophila melanogaster* and RNA interference, establishing the role of *An. gambiae* s.s. *CYP6Z1*, *CYP6Z2*, *CYP6Z3*, and *CYP6Z4* in conferring multiple- and cross-resistance towards pyrethroids, carbamates, organophosphates, and neonicotinoids, while increasing susceptibility to the pro-insecticide chlorfenapyr. These findings uncover the complex, multifaceted nature of metabolic resistance and provide valuable insights for the design and implementation of sustainable vector control and resistance management.

## 2. Materials and Methods

### 2.1. Mosquito samples

Larvae of *Anopheles gambiae s.s*. were collected between August 2022 and December 2023 in Mangoum, West Region of Cameroon (5°29′09.2″ N, 10°35′20.8″ E), a location previously described (Tepa et al., 2022). This site is characterized by intensive manual and mechanized agriculture, including cultivation of spices, vegetables, and cereals, providing an ideal breeding sites polluted with insecticides (Kengne-Ouafo et al., 2024). Briefly, breeding sites were randomly selected in each location, and larvae were collected using a 35 mL dipper. In parallel, indoor-resting adult female *An. gambiae s.s.* were collected in the early morning (06:00 to 08:00 a.m.) using battery-powered aspirators (John W. Hock, Gainesville, FL, USA). Larvae and adults were transported to the insectary of the Centre for Research in Infectious Diseases (CRID) in Yaoundé, and the larvae reared under standard insectary conditions and fed with Tetramin™ baby fish food (Tetra GmbH, Melle, Germany). Upon pupation and emergence, both field-collected (F₀) and laboratory-reared (F₁) adult mosquitoes were morphologically identified using the taxonomic keys (Gillies and De Meillon, 1968), and to species level using PCR. For all subsequent experiments, *An. gambiae s.s.* were used alongside their insecticide-susceptible laboratory strains, Kisumu.

### 2.2. Transcriptional profiling of CYP6Z genes from pyrethroid-resistant populations

Building on the previous RNAseq analysis conducted by Tepa et al. (2025), we reanalysed transcriptomic data following the pipeline developed in EasyRNAseq Repository (https://github.com/ArnaudTepa/EasyRNAseq) with a specific focus on the expression levels of *CYP6Z* genes, which are implicated in pyrethroid resistance in *Anopheles gambiae s.s.*, polymorphism detection and determination of population genetics metrics. The level of expression of *CYP6Z* genes family previously associated with pyrethroids resistance (Tepa et al., 2025) was validated by qRT-PCR, using three technical replicates of cDNA extracted from total RNA of three biological replicates each from the Resistant (R) and Control (C) *An. gambiae*, as well as susceptible laboratory colony, Kisimu (S). Briefly, 1 µg of the total RNA from each of the three biological replicates was used as the template for cDNA synthesis using Superscript III (Invitrogen, Carlsbad, CA, USA) with oligo-dT20 and RNase H, according to the manufacturer’s instructions. The qRT-PCR amplification was performed as described (Schmittgen and Livak, 2008) using the primers provided in Supplementary Table 1. The relative expression of *CYP6Z* gene was calculated as previously described (Tepa et al., 2022) by comparing expression in resistant, susceptible and control samples, after normalisation with the housekeeping genes ribosomal protein S7 (AGAP010592) and Elongation factor (AGAP000883). Significant differences were calculated using ANOVA with Dunnett’s post hoc test.

### 2.3. Polymorphism analysis of An. gambiae s.s. CYP6Z cDNA

To investigate patterns of genetic variability of *An. gambiae CYP6Z1, -Z2, -Z3* and *- Z4* and detect potential signatures of selection, the coding region of these genes were amplified from female of pyrethroid-resistant and lab susceptible mosquitoes and cloned. Total RNA from pools of 10 permethrin-resistant female *An. gambiae* s.s. from Mangoum and Kisumu lab susceptible strain were extracted using the PicoPure RNA isolation kit (Arcturus, Applied Biosystems, Foster City, California, USA). The purified RNA was used for cDNA synthesis using SuperScript III (Invitrogen) with oligo-dT20 and RNase H (New England Biolabs, Oxfordshire, UK). PCR amplification of *CYP6Z* genes was carried out using Phusion Hot Start II Taq polymerase (Thermo Fisher Scientific, MA, USA), following previously published protocols (Ibrahim et al., 2018). The following cycling parameters were used: 1 cycle at 98°C for 1 min; 35 cycles of 98°C for 10 s; 60°C for 30 s; and 72°C for 1 min and 20 s; and 1 cycle at 72°C for 10 min. The primers used are listed in Supplementary Table 1. PCR products were cleaned with a QIAquick® gel extraction Kit (QIAGEN, Hilden, Germany) and ligated into the pJET1.2/blunt cloning vector using the CloneJET PCR Cloning Kit (Thermo Fisher Scientifc, MA, USA). Ligated products were cloned into *E. coli DH5α*, plasmids miniprepped with the QIAprep® Spin Miniprep Kit (QIAGEN, Hilden, Germany) and sequenced on both strands using pJET1.2 specific primers.

To detect polymorphism and/or mutations potentially associated with the resistance phenotype, the above sequences were examined and manually edited using BioEdit version 7.2.5.0 (Hall, 1999). Population genetics parameters including number of haplotypes (h) and its diversity (H_d_), number of polymorphic sites (S) and nucleotide diversity (π) were computed using DnaSP 5.1 software (Librado and Rozas, 2009). Different haplotypes were compared by constructing a maximum likelihood phylogenetic tree using MEGAX (Kumar et al., 2018) and haplotype networks were constructed using the TCS program (Clement et al., 2000).

### 2.5. Homology modelling and docking simulations

To investigate the abilities of the *CYP6Z* genes to bind insecticides , homology models of the *An. gambiae s.s.* CYP6Z proteins selected from predominant alleles were developed using AlphaFold2 within the ColabFold open-source platform (Mirdita et al., 2022). Models for each allele were selected based on the highest average predicted local distance difference test (pLDDT) reliability scores for use in subsequent simulations. Ligand structures were sourced from the PubChem database (https://pubchem.ncbi.nlm.nih.gov/). The three-dimensional models of the proteins and ligands were prepared for docking using MGL Tools integrated within AutoDock (Kuntal et al., 2010). The heme ligand was retrieved from crystal structures that closely resembling our target enzymes, available in the Protein Data Bank (https://www.rcsb.org/), and was incorporated into the query model using PyMOL 3.0 (DeLano and Bromberg, 2004). The human microsomal cytochrome P450 *CYP3A4* (PDB 4D6Z) served as templates to predict the binding pocket of the heme (Yano et al., 2004) . Docking was performed using specific site docking with AutoDock Vina and active site defined as a cavity of 15 Å radius centred above the heme iron (Sánchez-Linares et al., 2012). The docking protocol was first optimized and validated by systematically adjusting pairs of scoring function parameters. This involved re-docking the native heme molecule into the 3D apo structure of CYP3A4 as well as into heme-mapped binding pockets of the predicted models, with the goal of accurately reproducing the original heme binding orientation (Supplementary Fig. 7).

Hundred (100) binding poses were obtained for each ligand for 1R-cis permethrin, deltamethrin, α-cypermethrin, bendiocarb, DDT, malathion, propoxur, chlorfenapyr, tralopyril, acetamiprid, clothianidin and imidacloprid (Irwin and Shoichet, 2005), which were sorted according to their binding free energy values in kcal/mol and the conformation of ligands in the active site of CYP6Z models (Salentin et al., 2015). Productive pose were define as poses with negative binding free energy and with distance from the iron heme range from 1.5 to 6.5 Å (Tatchou-Nebangwa et al., 2024). The conformations with the highest binding affinities were prioritized for an in-depth analysis of ligand-receptor interactions, with visualization facilitated by PyMOL 3.0 (DeLano and Bromberg, 2004).

### 2.6. Cloning and heterologous expression of recombinant CYP6Z3

Recombinant CYP6Z3 protein was expressed as previously described (Ibrahim et al., 2016). Briefly, expression plasmids were constructed (primers provided in Supplementary Table 1) by fusing cDNA fragment from a bacterial ompA+2 leader sequence with its downstream ala-pro linker to the NH_2_-terminus of the P450 cDNA, in frame with the P450 initiation codon (Pritchard et al., 1997). These constructs were digested with the *Bgl*II/*EcoR*I and *Xba*I restriction enzymes and ligated into pCW-ori+ expression vector, linearised with the same restriction enzymes, creating the constructs pB13::OMPA+2-CYP6Z3. The *E. coli JM109* cells were co-transformed with the above P450 constructs and a plasmid containing the *An. gambiae* cytochrome P450 reductase (pACYC-AgCPR). Cultured cells in LB medium were allowed to grow until reaching an optical density at 600nm of 0.7-0.8 before heme precursor δ-aminolevulinic acid (ALA), to a final concentration of 0.5 mM and isopropyl-1-thio-β-D-galactopyranoside (IPTG) to a final concentration of 1 mM were added. Membranes were isolated as done previously and P450 contents determined using spectral analysis (Omura and Sato, 1964), while CPR activities were done using established protocol (Guengerich et al., 2009). Purification of the recombinant CYP6Z3 protein was conducted as previously described (Ibrahim et al., 2015; Pritchard et al., 1997; Zelasko et al., 2013). Briefly, about 22h post induction, cells were harvested and spheroplasts were prepared and sonicated. The membrane fractions containing P450s were then isolated by ultracentrifugation at 50,000g and resuspended in TSE buffer (50 mM Tris, pH 7.6, 250 mM sucrose, 10% glycerol), and stored in -80°C following measurement of P450 contents.

### 2.7. In vitro metabolism assays with insecticides

Insecticide metabolism assay was carried out using previously described approaches (Tatchou-Nebangwa et al., 2024; Wamba et al., 2021). Briefly, 0.1µM of purified P450, 0.025 M potassium phosphate at pH 7.4, 0.25 mM MgCl_2_, 1 mM glucose-6-phosphate, 1 unit/mL glucose-6-phosphate dehydrogenase (G6PDH), 0.1 mM NADP was and 0.8 µM cytochrome b_5_ were added to 1.5ml tubes chilled on ice. The tubes were preincubated at 30°C and 1200 rpm for 5 min to activate membranes before 0.2 mM of insecticide to tested (permethrin, deltamethrin, α-cypermethrin, bendiocarb, clothianidin or fenitrothion) were added to the tubes, making a final volume of 200 ml. Tubes were shaken at 30°C and 1200 rpm for 120 min. All reactions were carried out in triplicate, with test reactions containing NADPH (NADPH+) and negative controls, lacking NADPH (NADPH-). After 2 h of incubation, 200 µl of ice-cold methanol (HPLC grade, Fisher Scientific) was added to the tubes to stop the reactions and the tubes were incubated for a further 5 min. The samples were then centrifuged at 16,000 rpm for 10 min at 4°C, and 150 μl of the supernatant was transferred into HPLC vials. The quantity of insecticide remaining in the samples was determined by reverse-phase HPLC (Agilent 1260 Infinity). For pyrethroids (permethrin, deltamethrin, and α-cypermethrin), detection was performed at 226 nm using an isocratic mobile phase of 90% methanol and 10% water. For bendiocarb, detection was set at 226 nm with an isocratic mobile phase of 60% acetonitrile and 40% water (Yunta et al., 2019). Pirimiphos-methyl and fenitrothion were monitored at 232 nm, with fenitrothion separated using an isocratic mobile phase of 80% methanol and 20% water (Ulusoy et al., 2020). All analyses were performed at 23°C with 1 mL/min flow rate on a 250 mm C18 column (Acclaim™ 120, Dionex), with injection volumes of 100 µl except for bendiocarb for which it was separated at 40°C. Metabolism was calculated as percentage depletion of insecticide (difference in the amount of insecticide left) between the test (NADPH+) and control (NADPH-). Student t-test was used for the estimation of significance levels (Kim, 2015).

### 2.8. Functional validation of the role of An. gambiae s.s. CYP6Z gene family in insecticide resistance using RNA interference

#### Double strand RNA synthesis and confirmation of CYP6Z genes knockdown using qRT-PCR

To investigate the impact of the knockdown of *CYP6Z1, -Z2, -Z3* and *-Z4* on. resistance, RNAi was performed using double-stranded RNAs synthesized from *CYP6Z* genes following the protocol we previously described (Kouamo et al., 2021). Each *CYP6Z* oligonucleotide primer was designed using specific cDNA of the corresponding genes and The T7 RNA polymerase promoter sequence, TAATACGACTCACTATAGGGAGA, was added to the 5′ end of all the primers (Supplementary Table 1). After amplification of the *CYP6Z* fragments by PCR from plasmid clones using KAPA Taq Kit (Kapa Biosystems, Wilmington, MA USA), each *CYP6Z* Double-stranded RNA (dsRNA) was synthesized using *in vitro* transcription MEGAscript® T7 Kit (Ambion Inc., Austin, TX, USA) and purified using MEGAclear columns (Ambion Inc., Austin, TX, USA). The purified products were concentrated by ethanol precipitation and the dsRNA was resuspended in nuclease-free water and stored at −20 °C. Successful creation of dsRNA was confirmed by running 3 μL of dsRNA-diluted products in 1.5% agarose gel in a Tris-acetate-EDTA (TAE) buffer.

Total RNA was extracted from 3 pools of 5 ds6Z-injected and non-injected mosquitoes using TRIzol reagent (Gibco BRL, Gaithersburg, MD, USA). cDNA from each of the three biological replicates was synthesized using the Super-Script III (Invitrogen, Carlsbad, CA, USA) with oligo-dT20 and RNase H, according to the manufacturer’s instructions, and used to assess the extent of RNAi by measuring levels of gene expression after injection by qRT-PCR following protocol previously described (Kouamo et al., 2021). A standard curve of each gene was generated using serial dilutions of cDNA to assess the knockdown efficiency after injection and the quantitative difference in the level of *CYP6Z* expression between injected and non-injected mosquitoes. The qPCR amplification was carried out in a MX3005 real-time PCR system using Brilliant III Ultra-Fast SYBR Green qPCR Master Mix (Agilent, Santa Clara, CA, USA). Briefly, 10 ng of cDNA from each sample was used as a template in a three-step program involving a denaturation at 95°C for 3 min followed by 40 cycles of 10 s at 95°C and 10 s at 60 °C and a last step of 1 min at 95°C, 30 s at 55°C and 30 s at 95°C. Using the 2^-ΔΔCT^ Livak procedure (Livak and Schmittgen, 2001), the relative expression and fold change of each target gene was calculated, comparing the expression in ds6Z-injected and non-injected samples after normalisation with the housekeeping genes S7 (AGAP010592) and Elongation factor (AGAP000883) as described above (section 2.2).

#### injection of mosquitoes and susceptibility bioassays

The dsRNA samples were injected into 2–3 d old F_1_ female mosquitoes from Mangoum followed by insecticide bioassays. A combination solution of double strand *ds6Z2+ds6Z1*, *ds6Z2+ds6Z3* and *ds6Z2+ds6Z4* were injected in the thorax of female mosquitoes using A Nano injector (Nanoinject; Drummond, Burton, OH, USA) as previously described (Blandin et al., 2002). Briefly, mosquitoes were anaesthetised with CO_2_ and injected with 69 nL of either aliquot of dual *dsCYP6Z2*+*6Z1*, *dsCYP6Z2*+*6Z3*, *dsCYP6Z2*+*6Z4* or dsGFP (control). After four days, four replicates of 20 mosquitoes for each treatment were exposed to permethrin (0.75%), α-cypermethrin (0.05%), deltamethrin (0.05%) and clothianidin (4%) for 1 h following the WHO testing protocol (WHO, 2016). To study the impact of the expression *CYP6Z* genes family on resistance intensity, double strand injected mosquitoes were also exposed to permethrin and α-cypermethrin 5X and 10X. After 1 h contact with insecticide, mosquitoes were transferred to holding tubes, supplemented with sugar and mortalities counted at 24 h after the exposure. The susceptibility test was performed in triplicates. Mosquitoes injected with *dsGFP* and those not injected were used as controls.

### 2.9. Investigation of the role of An. gambiae s.s. CYP6Z1, -Z3 and -Z4 in resistance using transgenic expression in Drosophila flies

#### Construction of transgenic flies

To assess whether overexpression of the *An. gambiae s.s. CYP6Z* genes alone could confer cross-resistance to different classes of insecticides, transgenic *Drosophila melanogaster* flies expressing *CYP6Z1*, *-Z3*, and *-Z4* were generated and tested with three pyrethroids (permethrin, deltamethrin and α-cypermethrin), bendiocarb (a carbamate), fenitrothion (an organophosphate), clothianidin (a neonicotinoid) and chlorfenapyr (a pyrrole) insecticides. The above three P450s were amplified using primers bearing *Bgl*II and *Xba*I restriction sites (Supplementary Table 1). The PCR amplicons were purified and cloned into pJET1.2 vector and miniprepped and predominant allele were digested from pJET1.2 vector Plasmids using the *Bgl*II and *Xba*I enzymes (Fermentas, Burlington, Ontario, Canada) and ligated into the pUASattB vector, pre-digested with the same restriction enzymes, and transformed into *E. coli DHα* cells (Invitrogen, Inchinnan Business Park, Paisley, UK) as previously described (Riveron et al., 2014). The recombinant constructs pUAS::CYP6Z1, pUAS::CYP6Z3, pUAS:CYP:6Z4 and pUAS::CYP6Z3 were injected into the germ-line of *D. melanogaster* carrying the attP40 docking site on chromosome 2 (y1 w67c23; P (CaryP) attP40,1;2) using the PhiC31 system (Markstein et al., 2008). Injection of flies and balancing were carried out by Cambridge Fly Facility (https://www.flyfacility.gen.cam.ac.uk/). Ubiquitous expression of the above constructs was obtained in the flies by crossing them with the driver line, Act5C-GAL4 strain (y1 w*; P (Act5C-GAL4-w) E1/CyO,1;2) (Bloomington Stock Center, IN, USA). Flies without UAS insert (white eyes) were also crossed with the Act5C-GAL4 line to create the control line. The expression of the recombinant CYP6Z1, -Z3, and -Z4 in the experimental transgenic flies was confirmed by semi-quantitative PCR using RNA extracted from tree pools of 10 F_1_ flies obtain from crossing of each transgenic line with Gal4-Actin 2 as previously described (Djoko Tagne et al., 2024; Kouamo et al., 2025).

#### Insecticides contact bioassays

The F_1_ progenies (3-4 d old females) overexpressing the above P450s were tested with insecticides using previously described protocols (Kouamo et al., 2025; Riveron et al., 2014; Tatchou-Nebangwa et al., 2024). The experimental flies and the control files were exposed for 24 h to 4% permethrin, 0.2% deltamethrin, 0.0007% α-cypermethrin, 4% DDT, 0.007% bendiocarb, 0.01% propoxur), 0.2% malathion, 0.2% pirimiphos-methyl, 2% fenitrothion, 50 µg/ml clothianidin, 40 µg/ml imidacloprid, 10 µg/ml acetamiprid and 10 µg/ml chlorfenapyr. Minimum of five replicates of 20 to 25 flies each were used for the bioassays, and the knockdown were scored after 1 h, 2 h, 3 h, 6 h, 12 h, and 24 h. Mortality and knockdown rates were compared between experimental and control groups using Student’s t-test.

## 3. Results

### 3.1. Transcriptional profiling of CYP6Z in pyrethroid-resistant An. gambiae s.s. populations

The RNA-seq analysis of *An. gambiae s.s. CYP6Z* locus reveals marked differential expression among resistant, control, and susceptible groups. Notably, *CYP6Z1, CYP6Z2*, and *CYP6Z3* genes exhibit varying levels of overexpression in resistant mosquitoes compared to susceptible and/or control groups, whereas *CYP6Z4* exhibited no notable upregulation across any comparison, suggesting a limited role in resistance (Supplementary Table 2).

*CYP6Z3* displayed the highest expression, with fold change of 41.4 (R-S) and 63.8 (C-S) indicating strong constitutive expression in resistant populations. Similarly, *CYP6Z2* was overexpressed, with fold changes of 17.2 (R-S) and 29.0 (C-S). Notably, *CYP6Z1* exhibited moderate but significant overexpression (6.0 in R-S and 12.3 in C-S), with substantial read counts (up to 19 358 in resistant samples). In contrast, the expression of *CYP6Z4* remained consistently low across all conditions (read counts between 40–86, no significant fold change).

### 3.2. Confirmation CYP6Z genes overexpression using qPCR

A qPCR analysis confirmed that *CYP6Z1*, *CYP6Z2*, and *CYP6Z3* were overexpressed in Mangoum population. However, the level of expression was not significantly different between permethrin-resistant populations and the unexposed controls (Figure 1). Specifically, *CYP6Z3* showed the highest expression, with fold change of 10.24 ± 4.15 in permethrin resistant, and 8.77 ± 3.29 in control mosquitoes. Similarly, *CYP6Z1* were upregulated in control (10.24 ± 4.29) and resistant mosquitoes (7.76 ± 1.28). In addition, *CYP6Z2* and *CYP6Z4* were also expressed in the Mangoum population, though its expression did not differ significantly between the resistant group (5.00 ± 1.66 and 3.74 ± 0.91, respectively) and the control group (3.64 ± 0.69 and 3.95 ± 1.01; p > 0.05, respectively).

**Figure 1:**
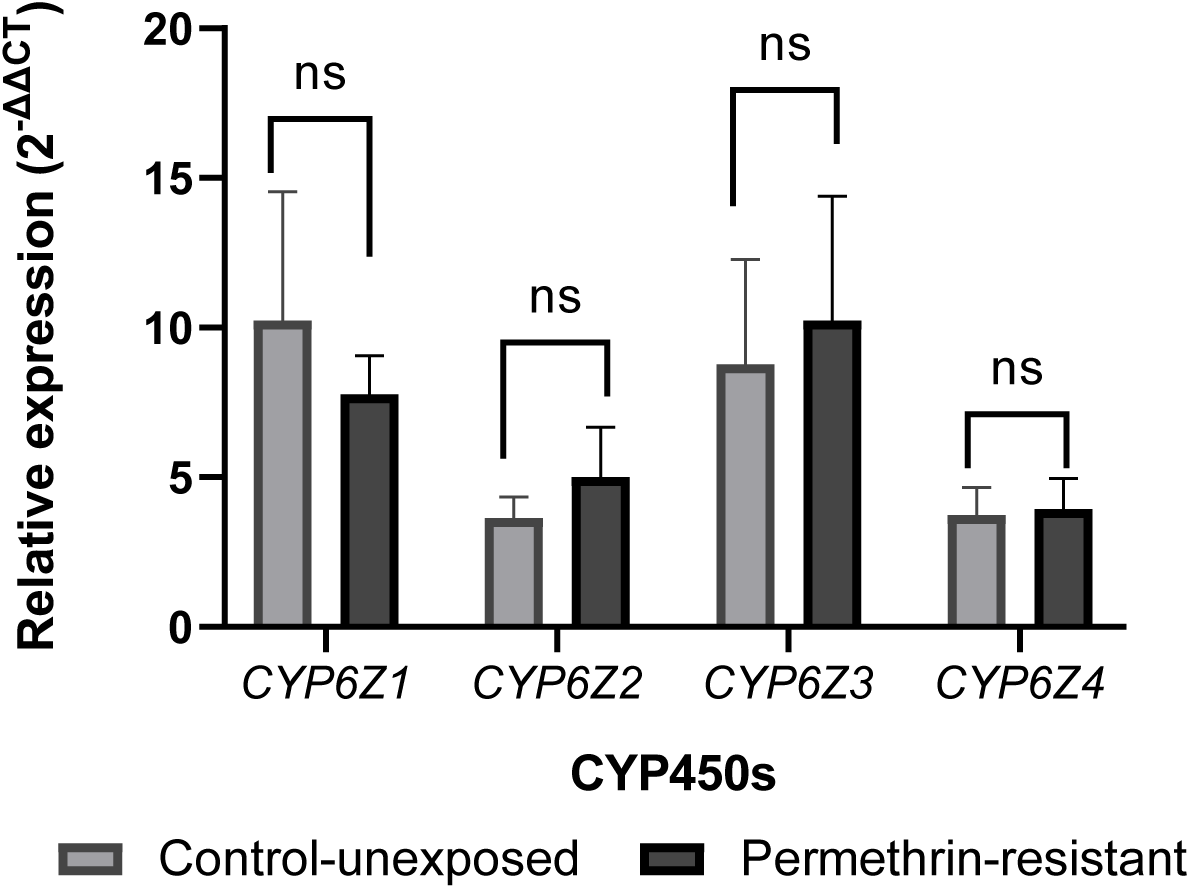
Differential expression profile of *CYP6Z* genes in permethrin-resistant *An. gambiae s.s.* field populations. Bar plots showing relative gene expression (2-^ΔΔCT^) of *CYP6Z1*, *CYP6Z2*, *CYP6Z3,* and *CYP6Z4* in permethrin-resistant versus control-unexposed mosquitoes. Significant overexpression was observed for *CYP6Z1*, *CYP6Z2*, and *CYP6Z3*, while *CYP6Z4* showed no significant change. ***p < 0.001, ns = not significant.

### 3.4. Genetic diversity analysis of An. gambiae s.s. CYP6Z genes

*Transcriptome-wide analysis of diversity and selective sweep across An. gambiae s.s. CYP6Z genes*

Using transcriptome derived single-nucleotide polymorphism (SNPs) analyzed at the gene level, we quantified population differentiation and neutrality across *CYP6Z* gene family (Supplementary Fig. 1). Gene wise F_ST_ was elevated in Mangoum permethrin resistant mosquitoes (Res.) vs Kisumu (Sus.), of 0.74 for *CYP6Z4*, 0.50 for *CYP6Z3* and 0.47 for *CYP6Z2* in R–S, with *CYP6Z1* at 0.38; dose response differentiation was detectable but lower for *CYP6Z1-3* [10×–1× F_ST_ = 0.1336 (*CYP6Z3*), 0.0332 (*CYP6Z2*), 0.0824 (*CYP6Z1*)] compared to *CYP6Z4* (0.76) (Supplementary Fig.1, A–D). Neutrality metrics supported localized selective sweeps, with negative Tajima’s D in Mangoum population, of −3.37 (*CYP6Z3*), −2.08 (*CYP6Z2*), and −1.23 (*CYP6Z1*), and reduced nucleotide diversity (π) relative to the susceptible strain (e.g., *CYP6Z3* π_field = 6.08 × 10⁻³ vs π_susceptible = 0; *CYP6Z2* π_field = 8.32 × 10⁻³ vs π_susceptible = 2.19 × 10⁻³; *CYP6Z1* π_field = 1.06 × 10⁻² vs π_susceptible = 6.29 × 10⁻³), consistent with recent selection acting within the *CYP6Z* cluster (Supplementary Fig.1, E–F).

Variant-level association analyses revealed strong selection signals for nonsynonymous SNPs within the *CYP6Z* cluster that were absent or very low in Kisumu, but highly enriched in field populations (Supplementary Fig.1.G). Within *CYP6Z1*, Ser279Asn (836 G > A) and Ser278Asn (833 G > A) mutations were observed, with highly significant and contrasts frequencies (26.32-40.12%) in field resistant mosquitoes, compared to Kisumu, in which they are completely absent (0%, p < 0.001). For *CYP6Z2*, the frequency of Ser201Thr (601 T > A) mutation increased significantly, from 26.51% in Kisumu to 64.23% in unexposed Mangoum and 50.24% at 10× permethrin (p < 0.05). Similarly, Asp380Glu (1140 C > G) increased from 0% in Kisumu to 33.48% at 5× permethrin. These patterns indicate dose-response enrichment of resistance-linked alleles under escalating insecticide pressure. For *CYP6Z3*, the frequency of Glu28Gly (83 A > G) mutation increased from 48.65% in Kisumu to 87.58–93.93% in Mangoum, supporting strong allele enrichment, although p values were not available for all contrasts because of coverage filtering.

#### Investigation of the genetic diversity linked with increased resistance intensity associated with CYP6Z genes

To investigate whether genetic diversity within the *CYP6Z* gene family is associated with permethrin resistance and its increased resistance intensity, we performed haplotype network analyses using coding sequences from Mangoum exposed to increasing concentrations of permethrin (1×, 5×, and 10×), unexposed field controls, and Kisumu colony.

The analyses revealed substantial haplotype diversity across the field population, with 35 haplotypes identified for *CYP6Z1* (Supplementary Fig. 2), 34 for *CYP6Z2* (Supplementary Fig. 3), 23 for *CYP6Z3* (Supplementary Fig. 4), and 24 for *CYP6Z4* (Supplementary Fig. 5). Haplotypes from the Kisumu colony consistently clustered distantly with longer branch lengths, reflecting their genetic divergence. However, with the exception of *CYP6Z2* haplotypes H15 and H19, and *CYP6Z4* haplotypes H14 and H15; which were shared among permethrin-resistant groups, no major haplotype was exclusively associated with resistance or intense resistance. Moreover, many haplotypes were either unique to specific exposure categories or dispersed across both resistant and control groups, suggesting a lack of clear directional selection or haplotype fixation under increasing insecticidal pressure. These data suggest that resistance mechanism may not be associated with amino acid mutations in the coding regions of the *CYP6Z* genes.

### 3.5. Polymorphism analysis of CYP6Z1, -Z2, -Z3 and -Z4 cDNAs

Comparative analysis of cDNA sequences revealed varying levels of polymorphism across the *CYP6Z* genes sub-family between the insecticide-resistant Mangoum mosquitoes vs Kisumu (Table 1). Haplotype network and genetic diversity analyses reveal high polymorphism and strong population differentiation in *CYP6Z* genes, with clear contrasts between the field populations and the susceptible lab colonies.

**Table 1:**
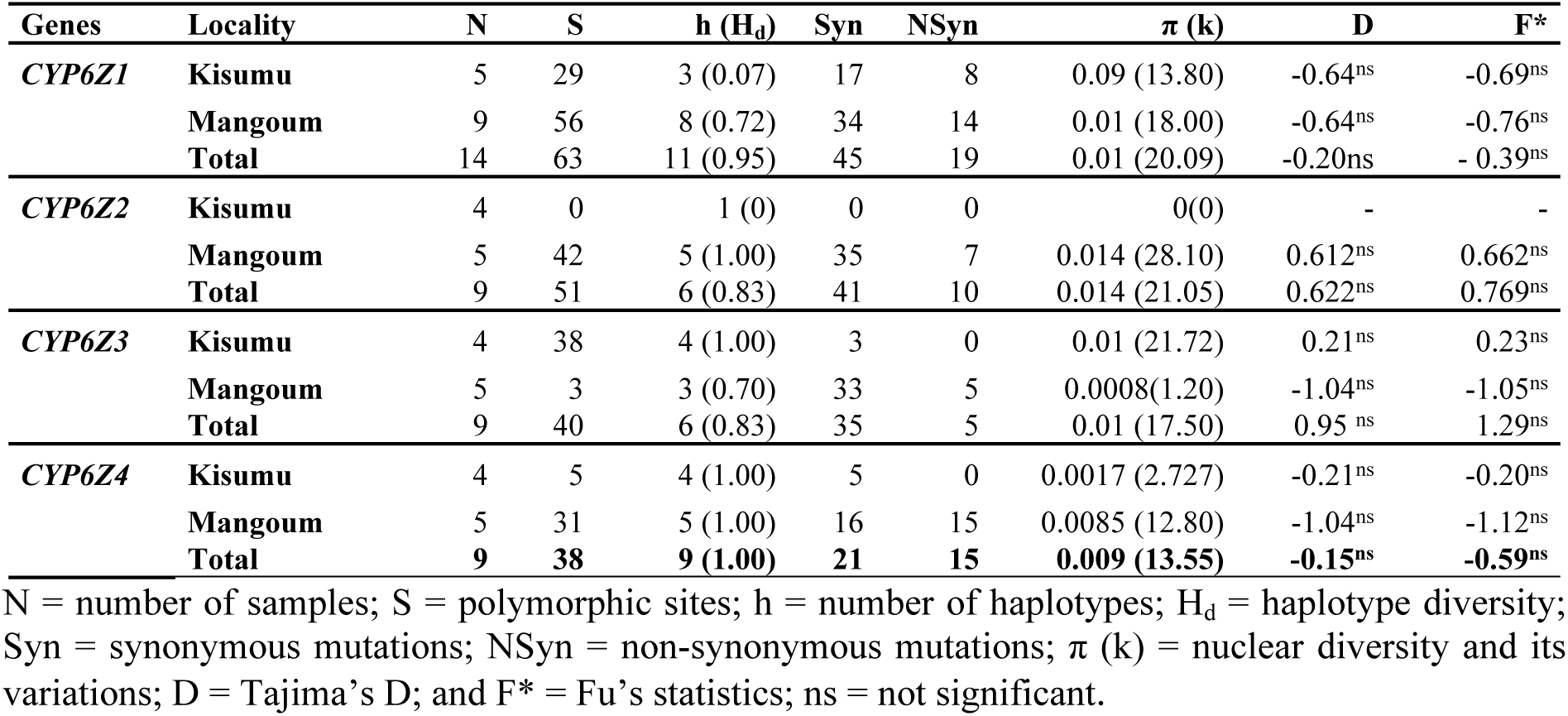
Summary statistics for polymorphism of An. *An. gambiae s.s. CYP6Z1, -2, -3 and -4* coding regions

*CYP6Z1* exhibited the highest genetic diversity H_d_ = 0.95 andπ = 0.01), with 11 haplotypes and 63 segregating sites, and a complex network structure. Key amino acid changes such as K151R-, S278N, S279N and Q377R which were detected in Mangoum were absent in Kisumu, suggesting that these variants may be linked to insecticide resistance (Supplementary Fig 6a, -b). In contract, *CYP6Z2* exhibited high diversity (H_d_ = 0.83, π = 0.014), with two amino acid changes H38Q (50%) and S201T-6Z2 (75%) present in Mangoum and absent in Kisumu (Figure 2). Similar pattern was observed for *CYP6Z3* with a high diversity (H_d_ = 0.83, π = 0.01), and few allelic variants: the T11A and E28G were fixed in Mangoum but present at 50% in Kisumu. *CYP6Z4* was found to be the most diverse (H_d_ = 1.00, π = 0.009), with two fixed amino acid changes V20I and A381P in Mangoum populations compared with Kisumu (Supplementary Fig 6c, -d).

**Figure 2:**
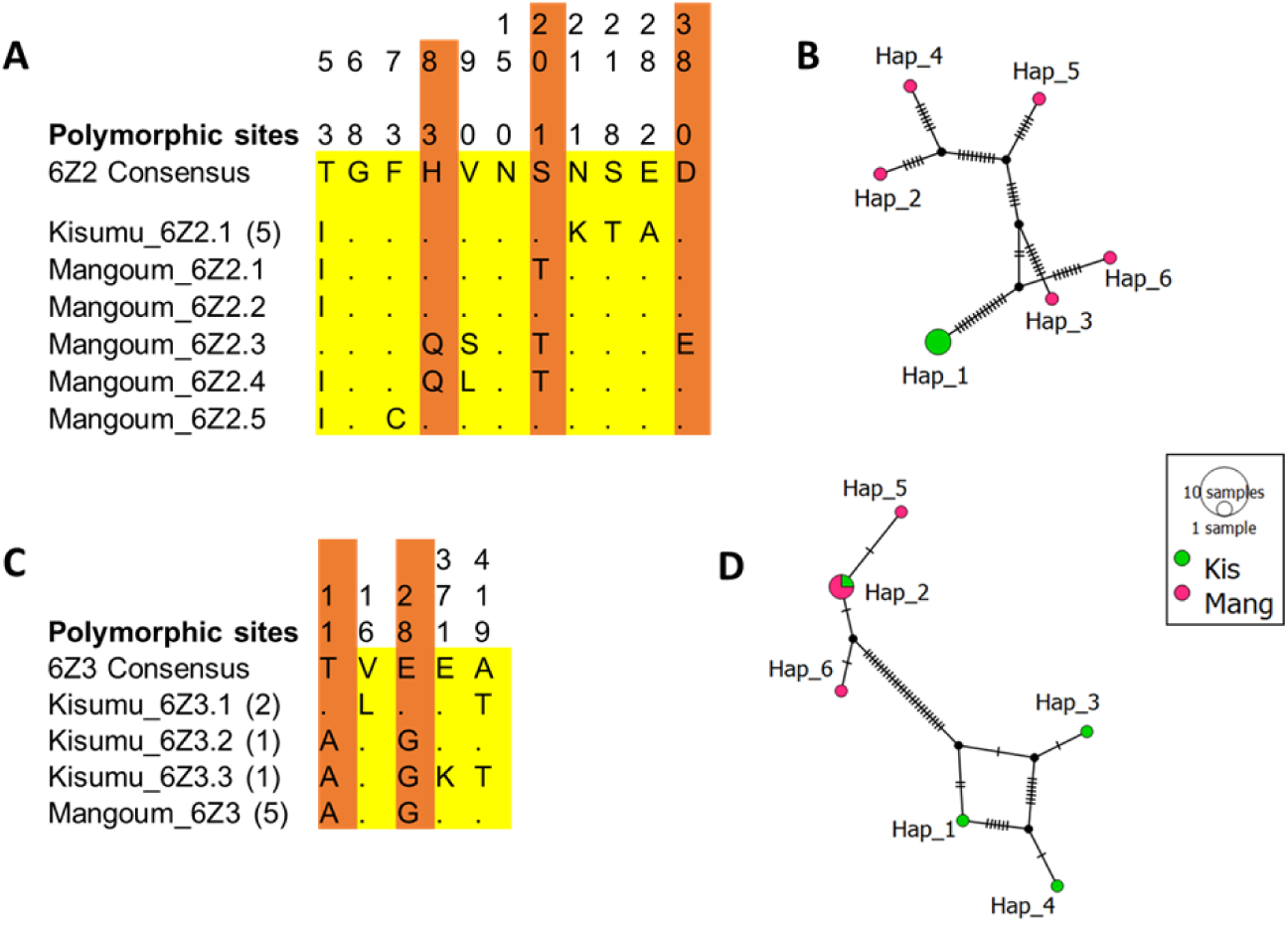
P**o**lymorphism **analysis of in *An. gambiae s.s. CYP6Z* genes coding regions.** (A) Comparative analysis of *CYP6Z2* amino acid changes between Mangoum *An. gambiae s.s.* and the Kisumu colony; (B) *An. gambiae s.s. CYP6Z2* haplotype diversity network showing that alleles from Mangoum do not share the same haplotype with Kisumu; (C) Comparative analysis of *CYP6Z3* amino acid changes between Mangoum *An. gambiae s.s.* and Kisumu sequences. (D) *An. gambiae* s.s. *CYP6Z3* haplotype diversity network showing that the dominant haplotype (H1) is shared between Mangoum and Kisumu.

### 3.6. Prediction of catalytic activities of CYP6Z genes using docking simulations

To predict the ability of the various *CYP6Z* genes to productively bind insecticides, 3D homology models of these P450s were created, and virtual insecticide structures docked into their active sites. The mean binding affinities for most of the various insecticides in the active sites of the four CYP6Z models showed no significant differences across (Supplementary Table 3). However, notable exceptions were observed: the Mangoum CYP6Z1 model exhibited a significantly lower binding energy with imidacloprid (-6.93 ± 0.31 kcal/mol) compared to the susceptible allele (-6.50 ± 0.33 kcal/mol, p = 0.003). Conversely, for CYP6Z2 model, the susceptible allele demonstrated higher affinity for α-cypermethrin (-8.35 ± 0.35 kcal/mol, p = 0.0004) and deltamethrin (-8.45 ± 0.33 kcal/mol, p = 0.0004) compared to the resistant allele (-8.88 ± 0.31 kcal/mol and -8.90 ± 0.40 kcal/mol, respectively). These findings indicate that, except for imidacloprid for CYP6Z1 and α-cypermethrin and deltamethrin for CYP6Z2, resistance-associated alleles do not exhibit significant differences in their predicted binding affinities toward insecticide ligands compared to susceptible allele from Kisumu.

Ligand-residue interaction analyses, combined with molecular docking simulations, provide a multidimensional view of insecticide metabolism across the two CYP6Z3 allelic models (Mangoum and Kisumu). LigPlot+ outputs for four insecticides classes (alpha-cypermethrin, malathion, clothianidin, and chlorfenapyr) revealed largely conserved hydrophobic contacts involving residues such as Phe^294^, Leu^295^, Ile^298^, Ala^299^, Leu^365^, and Val^477^, forming a lipophilic binding pocket in both alleles. In addition, hydrogen-bonding interactions with residues including Asn^113^, Ser^235^, and Thr^218^ were consistently predicted near the catalytic heme group across insecticides (Supplementary Fig. 7-8), although subtle differences in bonding patterns were observed between alleles and compounds. Permethrin adopts a productive pose in the CYP6Z1 and CYP6Z3 binding pockets, with the trans-methyl group oriented toward the heme at 3.5Å and 3.4Å, respectively (Supplementary Fig. 9). For type II pyrethroids, α-cypermethrin docked at 6.2Å in CYP6Z1 model (Supplementary Fig. 10) and 3.7Å in CYP6Z3 (Figure 3), while the C-4’ of the phenoxybenzyl ring of deltamethrin was positioned above the heme at 5.9Å for CYP6Z1 and 6.2Å for CYP6Z3, with the phenoxy ring favoring potential ring hydroxylation. For DDT, CYP6Z1 and CYP6Z3 bound with the trichloromethyl group of DDT positioned at a distance of 6Å and 4.6Å respectively from the heme. Among carbamates, propoxur docked productively with the C-5 of the phenyl ring pointing toward the heme iron at 3.4Å (CYP6Z3), whereas bendiocarb docked towards the heme at a distance above 6Å away from the C-4 phenyl ring of CYP6Z1 and CYP6Z3, for possible ring hydroxylation to 4-hydroxybendiocarb. For organophosphates, malathion was oriented closest to the heme iron at 3.4Å in CYP6Z3, suggesting highly favorable intermolecular interactions.

**Figure 3:**
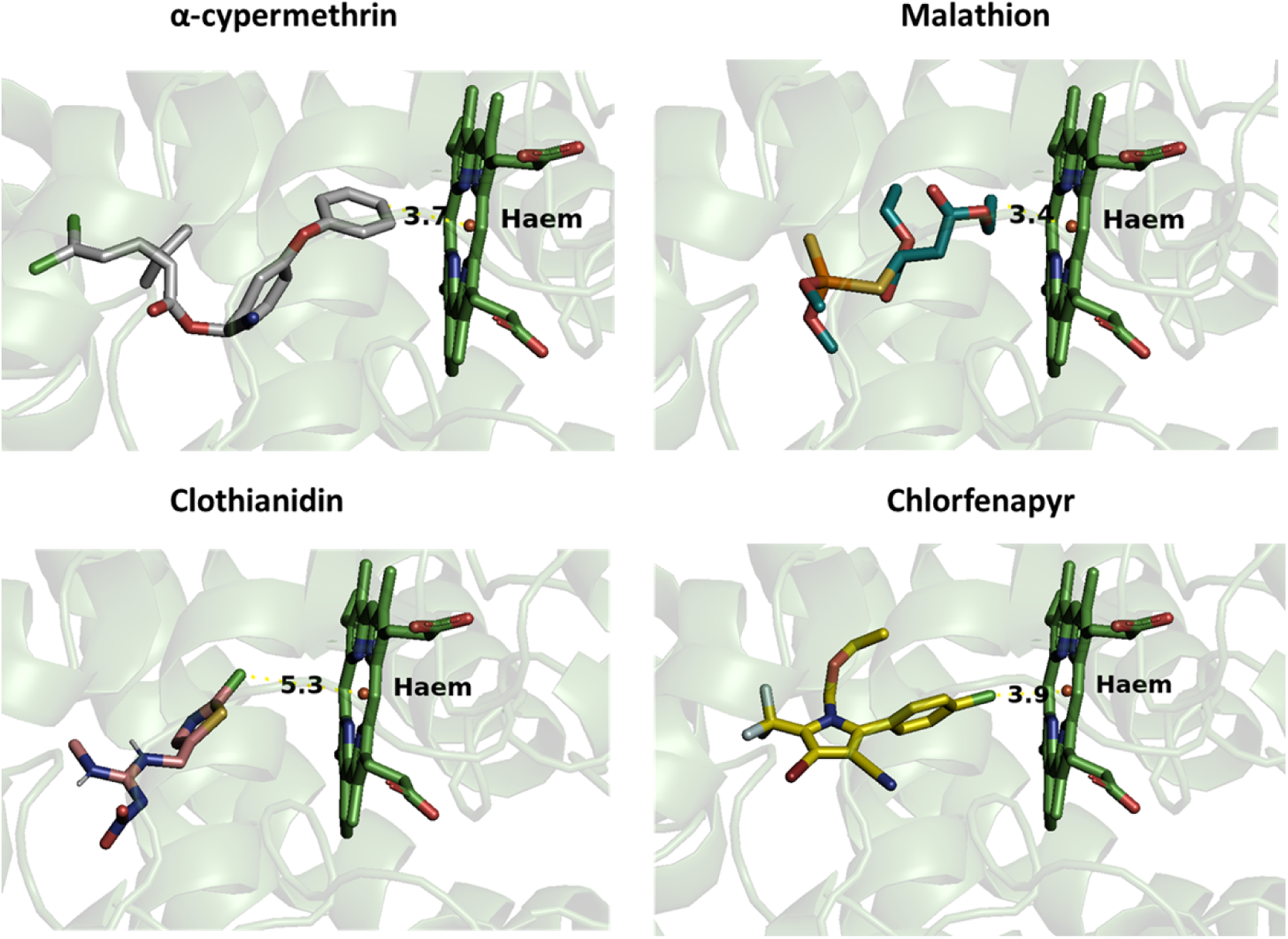
C**o**nformation **of various insecticides classes in *An. gambiae s.s*. *CYP6Z3* binding pocket.** Heme atoms are in stick format and colored by atom. Distances between possible sites of metabolism and heme iron are annotated in Angstrom.

Regarding neonicotinoids, the docking analysis indicated productive binding within the active site of CYP6Z3 model. Clothianidin was positioned at 5.3Å from the catalytic heme iron, suggesting a favorable orientation for oxidative metabolism. Imidacloprid was oriented with the nitrogen atoms of its heterocyclic moiety at 3.7Å from the heme, while acetamiprid docked at 4.4Å. These orientations suggest that CYP6Z3 could catalyze detoxification of imidacloprid via hydroxylation or N-demethylation. Additionally, CYP6Z3 docked productively with the ethoxymethyl group of chlorfenapyr and tralopyril at 3.9Å and 4.3Å, respectively, indicating a favorable orientation for the critical bioactivation of chlorfenapyr.

These structural models collectively highlight the ability of the CYP6Z protein family binding pocket to accommodate a wide variety of insecticide structures, each adopting a distinct orientation and distance relative to the catalytic heme. The diversity in binding modes and distances underscores the structural flexibility of *An. gambiae* s.s. CYP6Z1 and CYP6Z3, supporting their capacity to interact with and potentially metabolize multiple insecticide classes.

### 3.7. Recombinant CYP6Z3 metabolizes insecticides from several classes

To validate the ability of recombinant CYP6Z3 protein to detoxify insecticides, a HPLC-based metabolic assay was conducted. The recombinant CYP6Z3 exhibited the highest metabolic activity toward deltamethrin and permethrin with percentage depletions of 59.26% ± 3.88 and 52.98% ± 3.91, respectively (Figure 4A). Moderate depletion was observed for α-cypermethrin (37.14% ± 1.45), as well as for pirimiphos-methyl (36.00% ± 2.11). Lower depletions were observed for fenitrothion (23.00% ± 1.87), bendiocarb (17.00% ± 0.92) and propoxur (25.00% ± 3.14).

**Figure 4:**
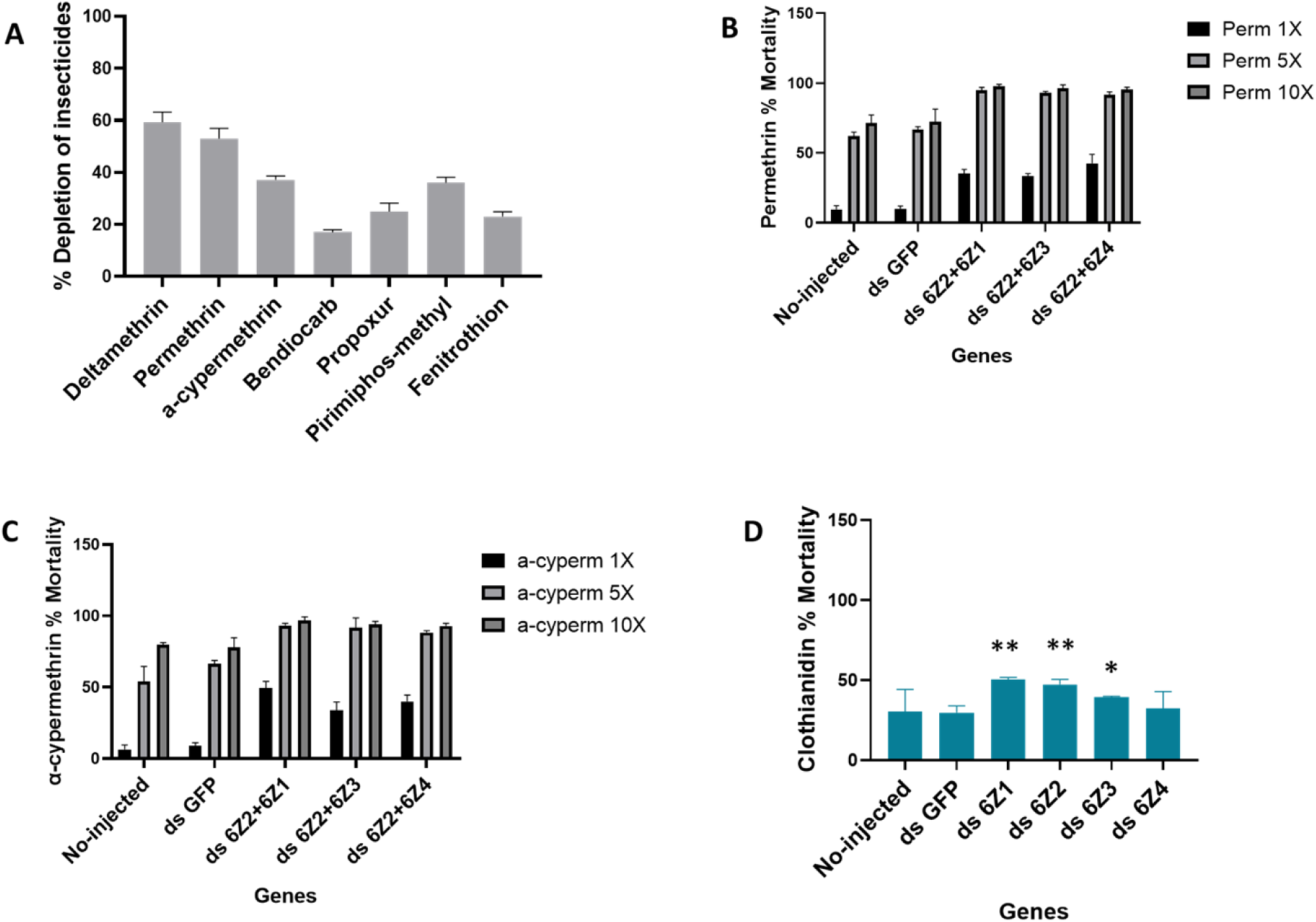
F**u**nctional **validation of the role of *An. gambiae s.s. CYP6Z* genes in conferring resistance.** (A) Percentage depletion of insecticides by recombinant CYP6Z3. Bars represent mean insecticide depletion (%) ± SEM from three independent experiments relative to negative control (no NADPH). (B), (C) and (D). Mortality rates of mosquitoes following injection of ds*CYP6Z* genes (RNAi), for permethrin, α-cypermethrin and clothianidin, respectively. * = p < 0.05, ** = p < 0.01, and *** = p < 0.001.

### 3.8. Knockdown of An. gambiae s.s. CYP6Z genes restores insecticide susceptibility

Silencing of the *CYP6Z* genes using RNAi confirmed the role of these P450s in insecticide resistance in Mangoum *An. gambiae* s.s., with mortalities increasing significantly in mosquitoes injected with dsCYP6Z combinations compared with the control group injected with dsGFP, and the non-injected mosquitoes. The reduced expression of *CYP6Z* in injected mosquitoes compared to the control, was confirmed by quantitative PCR (Supplementary Fig. 11). For 1X permethrin, mortality after 24 h of exposure were significantly higher in mosquitoes treated with dsCYP6Z2*+*CYP6Z1 (35.1% ± 3.2; P < 0.001), dsCYP6Z2*+*CYP6Z3 (33.1% ± 2.0; P < 0.01) and dsCY6Z2*+*CYP6Z4 (42.5% ± 6.4; P < 0.001) compared to the mortality observed in the control groups: dsGFP (10.1% ± 1.7) and non-injected (9.5% ± 2.6) mosquitoes (Figure 4B). No difference in mortality rate was found between mosquitoes injected with dsGFP and the non-injected mosquitoes, showing that the injection process did not affect fitness of the mosquitoes. For 5X permethrin, increased susceptibility was observed compared to mortality from 1X permethrin exposure. Mosquitoes injected with dsCYP6Z2+CYPZ1 (95.1% ± 1.8; P < 0.001), dsCYP6Z2+CYPZ3 (92.8% ± 1.1; P < 0.001), and dsCYP6Z2+CYPZ4 (91.6% ± 2.1; P < 0.01) were significantly more susceptible to 5X permethrin than control mosquitoes injected with dsGFP (66.5% ± 2.3) or non-injected (61.9% ± 3.1). A similar trend was observed from 10X permethrin exposure, where injected mosquitoes showed even higher susceptibilities. Specifically, dsCYP6Z2+CYPZ1 (97.7% ± 1.2; P < 0.001), dsCYP6Z2+CYPZ3 (96.3% ± 2.1; P < 0.001), and dsCYP6Z2+CYPZ4 (95.4% ± 1.6; P < 0.01) all displayed significantly higher mortalities compared to dsGFP-injected controls (72.2% ± 9.1), and non-injected mosquitoes (71.5% ± 5.6). In general, increased susceptibility was observed with higher insecticide concentrations, as mosquitoes exhibited greater mortality following exposure to 10X and 5X permethrin compared to 1X permethrin.

For α-cypermethrin, a significantly higher mortality at 24 h post exposure to 1X permethrin was observed among mosquitoes treated with dsCYP6Z2+CYPZ1 (49.2% ± 4.8; P < 0.001), dsCYP6Z2+CYPZ3 (33.1% ± 5.0; P < 0.01), and dsCYP6Z2+CYPZ4 (40.5% ± 4.5; P < 0.001), compared to control groups injected with dsGFP (8.9% ± 2.1), or non-injected mosquitoes (6.4% ± 3.1) (Figure 4C). As in the case of permethrin, a similar trend was observed when mosquitoes injected with combinations of *dsCYP6Z* were exposed to 5X and 10X α-cypermethrin. Notably, mosquitoes injected with dual dsCYP6Z2+CYP6Z1, dsCYP6Z2+CYP6Z3, and dsCYP6Z2+CYP6Z4 exhibited greater susceptibility after 5X and 10x α-cypermethrin exposure compared with mosquitoes exposed to 1X permethrin. Mosquitoes injected with dsCYP6Z2+CYPZ1 (93.34% ± 1.4; P < 0.001), dsCYP6Z2+CYPZ3 (91.68% ± 6.8; P < 0.001), and dsCYPZ2+CYPZ4 (88.31% ± 1.36; P < 0.01) displayed significantly increased susceptibility to 5X α-cypermethrin at compared to both dsGFP-injected controls (63.2% ± 2.3) and non-injected mosquitoes (54.11% ± 10.4). Similar pattern was observed following exposure to 10X α-cypermethrin, where mosquitoes injected with dsCYP6Z2+CYPZ1 (96.80% ± 2.5; P < 0.001), dsCYP6Z2+CYPZ3 (93.98% ± 2.2; P < 0.001), and dsCYP6Z2+CYPZ4 (92.96% ± 1.89; P < 0.001) were significantly more susceptible than the dsGFP-injected group (77.91% ± 6.8) and non-injected controls (79.88% ± 1.45). In the case of deltamethrin exposure, mortality rates recorded 24 h post-treatment were significantly higher in mosquitoes injected with dsCYP6Z2+CYPZ1 (38.6% ± 1.4; P < 0.001), dsCYP6Z2+CYPZ3 (30.1% ± 4.2; P < 0.001), and dsCYP6Z2+CYPZ4 (46.2% ± 3.4; P < 0.001), compared to the control groups: dsGFP (6.2% ± 2.3) and non-injected mosquitoes (5.06% ± 2.0) (Supplementary Fig. 12). However, the restoration of susceptibility after deltamethrin exposure was comparatively lower than what was seen with permethrin and α-cypermethrin.

A contrasting pattern of increased neonicotinoid susceptibility was observed following knockdown of *CYP6Z*. For example, a higher mortality rate was obtained with mosquitoes injected with dsCYP6Z1 (50.2% ± 1.4; P < 0.001), dsCYP6Z2 (47.2% ± 3.1; P < 0.001), dsCYP6Z3 (39.1% ± 3.2; P < 0.05) compared to the controls groups injected with dsGFP (29.6% ± 4.1) and the non-injected (29.6% ± 4.2), except for ds6Z4 injected files, which exhibited comparable mortalities (32.4% ± 5.3; P > 0.05) to the control groups (Figure 4D).

### 3.9. Recombinant CYP6Z genes confer resistance to transgenic Drosophila flies

To determine if overexpression of *CYP6Z* genes can alone confer cross-resistance to different classes of insecticides, transgenic *D. melanogaster* expressing *CYP6Z1, -Z2, -Z3* and -Z*4* were generated. After confirming expression in the transgenic flies and its absence in the controls using qPCR (Supplementary Fig.13), the flies were exposed to insecticides from six different classes: pyrethroids, carbamates, organophosphates, neonicotinoids, pyrroles, and organochlorines.

#### 3.9.1. CYP6Z genes confer pyrethroid and DDT resistance in transgenic flies

Transgenic flies expressing *CYP6Z1* were significantly more permethrin resistant, with mortalities of 4%, 9%, 18%, and 48% after 3h, 6h, 12h, and 24h, respectively (p<0.01), compared to control flies (25%, 39%, 53%, and 85% after 3h, 6h, 12h, and 24h, respectively) (Figure 5A). For *CYP6Z3* flies, significantly higher resistance was also observed with mortalities of 4%, 12%, 18%, and 41% after 3h, 6h, 12h, and 24h, respectively (p<0.01), compared with the control flies. Similar pattern was observed with *CYP6Z4* flies with permethrin mortalities of 3%, 10%, 22%, and 27% after 3h, 6h, 12h, and 24h, respectively (p<0.001) compared with control flies. Lower mortality was observed with transgenic flies expressing *CYP6Z1* 28%, 36% and 53% after 6, 12 and 24 h, respectively (p<0.01) upon α-cypermethrin exposure compared to control flies (47%, 75% and 94% after 6h, 12h and 24h, respectively). For *CYP6Z3* flies significantly higher α-cypermethrin resistance was also observed with mortality of 25%, 26% and 77% after 6, 12 and 24 h, respectively, (p<0.01) compared with the control above. Comparable trend was observed with *CYP6Z4* flies being more resistant (24%, 29% and 59% after 6, 12 and 24 h, respectively, (p<0.001)), compared to the control (Figure 5B). For deltamethrin, significantly low mortalities were obtained in transgenic flies expressing *CYP6Z1* at 24%, 41%, and 68% after 6h, 12h, and 24h, respectively (p<0.01), compared with the control (47%, 75%, and 94% after 6h, 12h, and 24h, respectively). *CYP6Z3* also confer low mortalities, of 18%, 22%, and 68% after 6h, 12h, and 24h, respectively, (p<0.01) compared with the control flies. A consistent trend was observed, with *CYP6Z4* transgenic flies showing significantly lower mortality rates of 10%, 25%, and 40% at 3h, 6h, 12h, and 24h, respectively (p < 0.001), compared to the control flies (Figure 5C). Between experimental arms comparisons revealed that *CYP6Z4* conferred higher permethrin (27%; P < 0.05) and deltamethrin (40%; P < 0.05) resistance than *CYP6Z1* (Supplementary Fig. 14). This contrasts with α-cypermethrin, from which transgenic flies expressing *CYP6Z1* (53%; P < 0.001) and *CYP6Z3* (47.2% ± 3.1; P < 0.001) were more resistant after 24 h exposure than transgenic flies expressing C*YP6Z4*.

**Figure 5:**
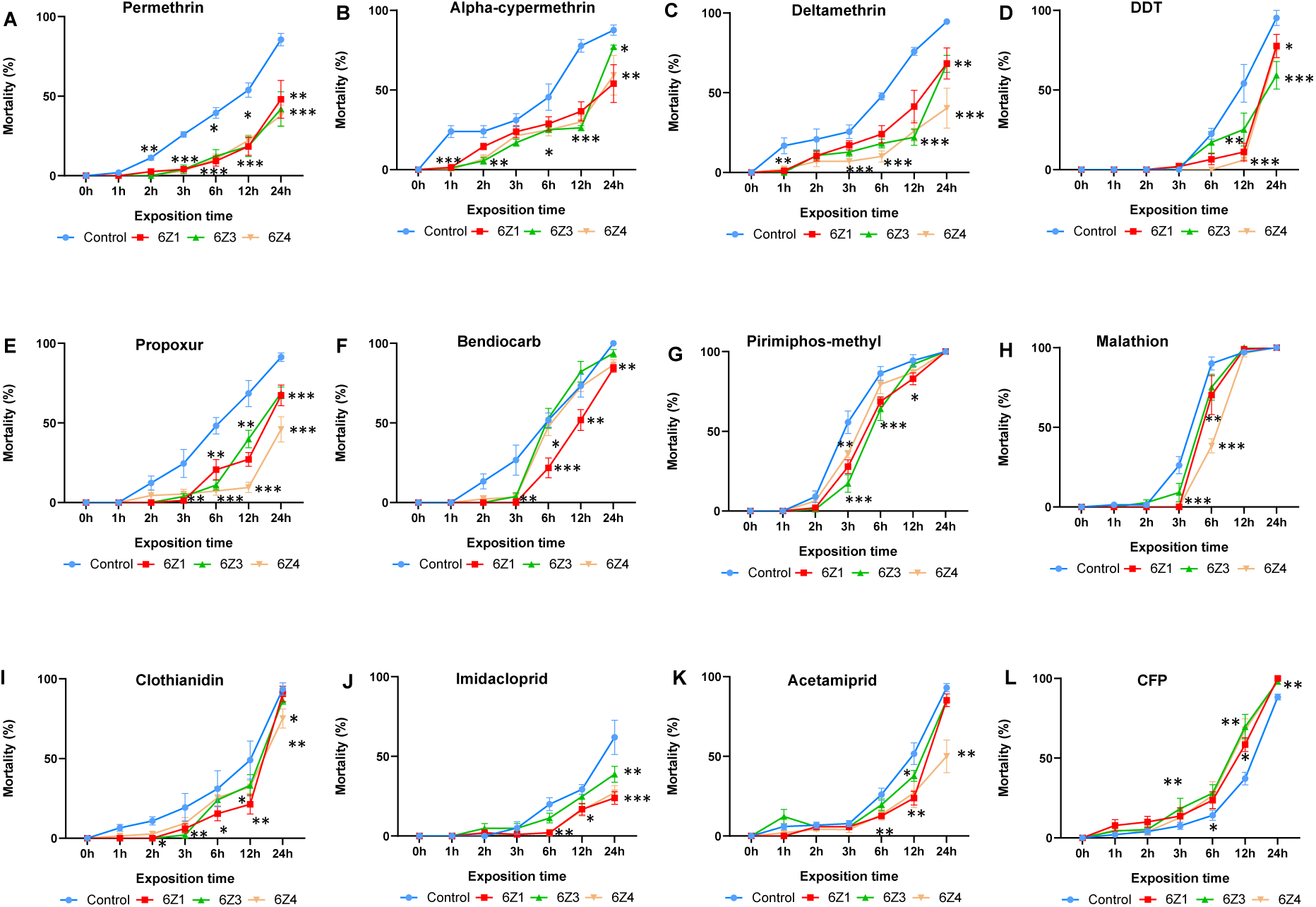
I**m**pact **of transgenic expression of *An. gambiae s.s. CYP6Z* genes on insecticide resistance.** Bioassay results showing mortality of transgenic flies expressing *CYP6Z* genes after exposure to different insecticide classes: (A) permethrin; (B) α-cypermethrin; (C) deltamethrin; (D) DDT; (E) propoxur; (F) bendiocarb; (G) pirimiphos-methyl; (H) malathion; (I) clothianidin; (J) imidacloprid; (K) acetamiprid; (L) chlorfenapyr. Statistical significance is indicated as follows: *p < 0.05; **p < 0.01; ***p < 0.001.

In addition, bioassays performed with DDT revealed that flies expressing *An. gambiae CYP6Z1* (11% and 77% after 6h and 24h, respectively), *CYP6Z3* (25% and 59% after 12h and 24h, respectively) and *CYP6Z4* (6% and 77% after 12 and 24h, respectively) were more resistant to DDT compared to the control (54% and 95% after 12h and 24h, respectively) (Figure 5D).

#### 3.9.2. CYP6Z genes confer carbamate resistance in transgenic flies

Recombinant *CYP6Z* genes confer propoxur resistance, with significantly lower mortalities of 20%, 27% and 60% after 6h, observed at 12h and 24h, respectively (p<0.01) for *CYP6Z1*; 10%, 39% and 68% after 6, 12 and 24 h, respectively (p<0.01) for *CYP6Z3*, and 7%, 9% and 47% after 6h, 12 and 24 h, respectively (p<0.001) for *CYP6Z4*, all compared to the higher mortalities obtained with control flies (48%, 68% and 91% after 6h, 12h and 24h, respectively) (Figure 5E). Transgenic flies expression *CYP6Z* genes were more resistant to bendiocarb, with significantly lower mortalities, of *CYP6Z1* (0%, 21%, 51% and 84% after 3h, 6h, 12 and 24 h, respectively (p<0.01) for *CYP6Z1*, 3% after 3h (p<0.01) for *CYP6Z3*, and 3% and 86% after 3h and 24h, respectively (p<0.01) for *CYP6Z4*, compare to the higher mortalities obtain from the control flies (26%, 51%, 73% and 100% after 3h, 6h, 12h and 24h, respectively) (Figure 5F). The *CYP6Z* genes confer higher resistance towards propoxur compared with bendiocarb.

#### 3.9.3. CYP6Z genes confer organophosphate resistance in transgenic flies

Transgenic flies expressing *CYP6Z1*, *-3* and *-4* were exposed to pirimiphos-methyl, malathion and fenitrothion. Flies expressing *CYP6Z* genes were more pirimiphos-methyl resistant, with a significant low mortality of 27%, 68% and 83% after 3h, 6h and 12h, respectively (p<0.05) for *CYP6Z1*, and 17%, 64% and 91% after 3h, 6h and 12h for *CYP6Z3*, respectively, compared to the control flies (55%, 86% and 94% after 3h, 6h and 12h, respectively (p<0.05) (Figure 5G). No differences in mortalities were observed in flies expressing *CYP6Z4* compared with control flies. The recombinant *CYP6Z* genes were observed not to confer malathion resistance in general (Figure 5H). However, subtle differences in susceptibilities were observed after 3h exposure. For example, at 3h flies expressing *CYP6Z* genes were more resistant, with low mortality of 0% (p<0.01) for CYP*6Z1*, 9% (p<0.01) for *CYP6Z3* and 0% (p<0.001) for *CYP6Z4*, compared to the low mortality of 26% observed with the control. Full susceptibility for all experimental and control flies was observed upon fenitrothion exposure (Supplementary Fig. 15).

#### 3.9.4. CYP6Z genes confer marginal neonicotinoid resistance in transgenic flies

Transgenic expression of *CYP6Z* genes increase clothianidin resistance, with low mortalities of 6%, 15%, 21% and 90% after 3h, 6h, 12h and 24h, respectively (p<0.05) for *CYP6Z1*, of 2%, 24%, 33% and 86% after 3h, 6h, 12h and 24h, respectively (p<0.01) for *CYP6Z3*, of 9%, 25%, 32% and 75% after 3h, 6h, 12h and 24h, respectively (p<0.05) for *CYP6Z4*, compared to the mortality observe with the control flies (19%, 31%, 49% and 93% after 3h, 6h, 12h and 24h), respectively (Figure 5I). A similar pattern was observed with imidacloprid with transgenic flies expressing *CYP6Z* genes exhibiting more resistance, with mortalities of 2%, 16% and 23% after 6h, 12h and 24h, respectively (p<0.05) for *CYP6Z1*, of 11%, 24% and 38% after 6h, 12h and 24h, respectively (p<0.05) for *CYP6Z2* and of 2%, 16% and 21% after 6h, 12h and 24h, respectively (p<0.05) for *CYP6Z4*, compared with mortality obtain with the control flies (19%, 29% and 61% after 6h, 12h and 24h), respectively (Figure 5J). Increased acetamiprid resistance was observed in transgenic flies, with mortalities of 12%, and 23% after 6h and 12h, respectively (p<0.05) for *CYP6Z1*, of 13%, 27% and 50% after 6h, 12h and 24h, respectively (p<0.05) for *CYP6Z4* compared with mortalities from the control flies (26%, 51%, and 93% after 6h, 12h, and 24h), respectively (Figure 5K). In contrast, no significant difference in mortalities was observed between flies expressing *An. CYP6Z3* and the control flies.

#### 3.9.5. CYP6Z genes increase chlorfenapyr susceptibility in transgenic flies

Bioassays performed with 10µg/ml chlorfenapyr (Figure 5L) revealed that the experimental flies expressing *CYP6P*Z gene were significantly more susceptible at 3h, 6h, 12h and 24h post-exposure than the control flies, with average of mortality at 24h of 100% for *CYP6PZ1* (P < 0.01), 98.1% ± 1.8 for *CYP6PZ3* (P < 0.01) and 98.8% ± 1.9 for *CYP6PZ4* (P < 0.01) compared to the control flies (88.3% ± 1.7). This indicates that the overexpression of these P450s alone increases the susceptibility to chlorfenapyr.

## 4. Discussion

Insecticide resistance remains a major challenge to malaria vector control, with increasing evidence of cross-resistance compromising the effectiveness of even newer chemical classes. Despite repeated associations of the *CYP6Z* gene family with resistance in *An. gambiae* mosquitoes, its precise functional contribution in cross-resistance to several insecticide classes and in chlorfenapyr bio-activation remain insufficiently understood. Addressing this knowledge gap, our study integrated transcriptomic analysis, genetic diversity, *in silico* structural characterization, *in vitro* and *in vivo* functional approaches to dissect the role of *An. gambiae* s.s. *CYP6Z1*, *CYP6Z2*, *CYP6Z3 and CYP6Z4*. Notably, we demonstrate for the first time the capacity of *CYP6Z* genes to bioactivate chlorfenapyr. These findings provide novel mechanistic insights essential for informing resistance management strategies and safeguarding current and future interventions.

### 4.1. Transcriptional overexpression of An. gambiae s.s. CYP6Z1, -Z2, -Z3 and -Z4: A consistent signature of insecticide resistance

The expression profiling of the *CYP6Z* gene family revealed this gene family is over-expressed in Cameroon, with particularly high levels observed in permethrin-resistant groups. The consistent and significant overexpression of *CYP6Z* genes in permethrin-resistant populations, especially *An. gambiae s.s. CYP6Z1*, *CYP6Z2*, and *CYP6Z3* strongly supports their role in the detoxification of permethrin. In the RNA-Seq dataset, *CYP6Z3* displayed the highest fold change (>39×) in resistant *An. gambiae* from Mangoum, highlighting it as a potential key player in constitutive resistance mechanisms. Similarly, *CYP6Z1* and *CYP6Z2* showed significant upregulation, corroborated by qPCR validation, where expression increased by over 25-fold in permethrin-resistant field populations relative to unexposed controls. This result is in line with previous studies where *CYP6Z1*, *CYP6Z2* and *CYP6Z3* genes have been functionally linked to insecticide metabolism in *An. gambiae* (Ingham et al., 2014; Nikou et al., 2003; Tepa et al., 2022; Tepa et al., 2025). This pattern is consistent with observations in West Africa, where *CYP6Z2* and *CYP6Z*3 were found to be highly overexpressed in resistant *An. coluzzii* from Burkina Faso and Nigeria respectively and were shown to metabolize pyrethroids and DDT (Kwiatkowska et al., 2013; Muhammad et al., 2021). Our finding supports previous finding highlighting the link between the overexpression of *CYP6Z* genes family in insecticide resistance observe in Sahelo-Sudanian *An. coluzzii* population, notably from northern Nigeria, southern Niger, central Chad and northern Cameroon (Ibrahim et al., 2023). In addition, the fact that *CYP6Z3* was both constitutively and inducible expressed in *An. coluzzii* and *An. gambiae* underlines the gene’s dual regulatory flexibility, making it an especially promising candidate for further functional validation (Müller et al., 2007). These findings collectively support the hypothesis that *CYP6Z* gene overexpression contributes significantly to metabolic detoxification pathways in pyrethroid-resistant malaria vectors.

### 4.2. An. gambiae s.s. CYP6Z genes family genetic variation as a contributor to resistance evolution

Beyond the critical role of altered gene expression, our comprehensive genomic and cDNA polymorphism analyses provide new insights into the genetic architecture and ongoing evolution of insecticide resistance within the *Anopheles gambiae CYP6Z* gene family. Transcriptome-wide analyses revealed robust indications of selection across the *CYP6Z* gene family in Mangoum, as demonstrated by elevated FST, considerably diminished nucleotide diversity, and markedly negative Tajima’s D patterns, which are conventionally linked to recent selective sweeps in insecticide-exposed Anopheles populations. (Clarkson et al., 2021; Weetman et al., 2018). Variant-level analyses further strengthened this interpretation by showing dose-dependent enrichment of nonsynonymous mutations across *CYP6Z1*, *CYP6Z2*, and CYP6Z3, which are rare or absent in the susceptible Kisumu strain but highly prevalent in field populations. These results collectively suggest that the *CYP6Z* genomic cluster is undergoing localised adaptive evolution in response to intense pyrethroid-based selection pressure. This is consistent with previous reports that metabolic gene clusters are key hotspots of resistance evolution in *An. gambiae* (Ingham and Nagi, 2024; Lucas et al., 2019). The genetic diversity and haplotype structure of the *An. gambiae CYP6Z* gene family reveal a complex evolutionary landscape shaped by insecticide selection pressures, where regulatory adaptation predominates over structural mutations in driving resistance evolution. With the high polymorphism detected across *CYP6Z1* in field populations, no major fixed resistance-associated alleles were identified, even under escalating permethrin exposure (1× to 10×). Key amino acid substitutions such as *CYP6Z1* S278N/S279N (42–56% frequency in field strains) and *CYP6Z3* V166A/E277V (26% frequency) emerged as transient markers of adaptation and their absence in lab-susceptible strains (0%) suggest that this allelic variation may contribute to the escalation of the insecticide resistance observed in the field. It supports the previous finding showing that Allelic variation can play an additional role in Anopheles P450-mediated resistance by modifying either enzyme catalytic activity or gene expression levels (Schuler and Berenbaum, 2013). Our finding is in line with previous study showing that key amino amino acid changes in *CYP6P9b* (V109I, D335E, N384S) between field resistant and lab susceptible strain drive bed net insecticide resistance in *An. funestus* (Ibrahim et al., 2015). Even *CYP6Z4*, despite its comparatively moderate role in overexpression in this context, presented high diversity and fixed amino acid changes (V20I and A381P) uniquely present in permethrin resistance population from Mangoum. These convergent accumulations of distinct amino acid substitutions across the *CYP6Z* gene family in resistant field populations directly demonstrate the adaptive fine-tuning of enzyme structure in response to selective pressure, thereby enhancing their capacity for insecticide detoxification (Kientega et al., 2024; Toé et al., 2015; Weetman et al., 2018).

### 4.3. Structural features of CYP6Z enzymes promote broad-spectrum insecticide cross-resistance through enhanced substrate binding and metabolism

The integration of metabolic assays and structural modeling provides compelling evidence that the structural features of CYP6Z proteins directly enable broad-spectrum insecticide cross-resistance in malaria vectors. The *in vitro* metabolic assay of *CYP6Z3* from *An. gambiae* demonstrates high depletion rates for pyrethroids with moderate activity toward carbamates and organophosphates, indicating a broad substrate range. The highest depletion rates were observed with deltamethrin and permethrin confirmed a strong detoxification potential for type II and type I pyrethroids, respectively. This aligns with phenotypic bioassays demonstrating cross-resistance to pyrethroid insecticides in *An. gambiae* populations overexpressing *CYP6Z3*. The moderate metabolic activity toward alpha-cypermethrin and organophosphates such as pirimiphos-methyl and fenitrothion further suggests that *CYP6Z3* may contribute to resistance across multiple insecticide classes, albeit with varying efficiency.

These functional results are supported by molecular docking and binding pocket analyses, which reveal that these insecticides are positioned within catalytically favorable distances (3.5–6.4 Å) from the heme iron in the CYP6Z enzymes family active site, facilitating efficient electron transfer and substrate oxidation (Liu et al., 2003). The close proximity and optimal orientation of diverse insecticides within the active site reflect the remarkable structural flexibility of *CYP6Z* enzymes, consistent with previous studies showing that *CYP6Z1*, *CYP6Z2*, and *CYP6Z3* confer metabolic resistance to multiple insecticide classes (Ibrahim et al., 2016; McLaughlin et al., 2008). This structural adaptability is further highlighted by the observed differences in binding energies and substrate positioning between resistant and susceptible alleles, supporting the notion that enhanced substrate binding and metabolism underlie cross-resistance phenotypes. Collectively, these findings reinforce the central role of *CYP6Z* enzymes as a cytochrome P450 in the metabolic detoxification of a wide spectrum of insecticides and underscore the molecular basis for cross-resistance in Anopheles vectors, as previously reported in field and laboratory studies (Moyes et al., 2020; Stevenson et al., 2011).

### 4.4. CYP6Z genes confer cross-resistance to several insecticide classes in transgenic flies

Transgenic expression of *CYP6Z* gene subfamily in *D. melanogaster* demonstrated that the overexpression of these gene plays a central role in conferring cross-resistance to multiple classes of insecticides, including pyrethroids, carbamates, neonicotinoids, and, to a lesser extent, organophosphates. In contrast, expression of these genes did not confer significant resistance to certain organophosphates, such as fenitrothion, which is in line with previous reports that P450-mediated resistance is often substrate-specific (Mitchell et al., 2012). Flies expressing *An. gambiae s.s. CYP6Z1*, *CYP6Z3*, and *CYP6Z4*, exhibited significantly lower mortality rates than controls when exposed to pyrethroid insecticides such as permethrin, deltamethrin, and α-cypermethrin. These findings are consistent with previous studies showing that overexpression of CYP6Z genes is a major mechanism underlying metabolic resistance in malaria vectors (Ibrahim et al., 2023; Ibrahim et al., 2016). Notably, the enhanced resistance was not limited to pyrethroids; transgenic flies also showed increased survival when exposed to carbamates (bendiocarb and propoxur), and new insecticide class such as neonicotinoids (clothianidin and imidacloprid), supporting the hypothesis that *CYP6Z* enzymes contribute to broad-spectrum metabolic resistance. This finding consistently shows in addition to pyrethroids, carbamate and DDT resistance previously demonstrated (Chiu et al., 2008; McLaughlin et al., 2008; Zouré et al., 2021) that overexpression of *CYP6Z* gene subfamily also play a role on cross-resistance to new insecticide such as clothianidin, imidacloprid and acetamiprid. Importantly, RNAi-mediated knockdown of dual *CYP6Z* genes in *An. gambiae* s.s. from Mangoum restored susceptibility to pyrethroids and neonicotinoids, confirming the causal role of these enzymes in resistance phenotypes. In addition, the RNAi result provide functional evidence that simultaneous knockdown of *CYP6Z2* in combination with either *CYP6Z1, CYP6Z3*, or *CYP6Z4* enhances the toxicity of α-cypermethrin, suggesting a synergistic contribution of these genes to cross-resistance to current deployed insecticides (Margaret et al., 2025; Minetti et al., 2020; Rahi et al., 2024; Tepa et al., 2025). Taken together, these results provide compelling functional evidence that *CYP6Z* gene subfamily members are key drivers of cross-insecticide resistance in malaria vectors, and highlight the need for resistance management strategies that target metabolic resistance mechanisms (Ibrahim et al., 2023; Mitchell et al., 2014). Collectively, these results underscore the importance of *CYP6Z* gene subfamily as a metabolic resistance factor, with implications for both the evolution of cross-resistance in field populations and the rational design of new vector control interventions targeting metabolic pathways.

### 4.5. CYP6Z genes enhance chlorfenapyr susceptibility: an encouraging finding for novel vector control strategies

A positive result from this study is the observed increase in mortality to chlorfenapyr in transgenic *D. melanogaster* expressing anopheles *CYP6Z* genes, especially *An. Gambiae s.s. CYP6Z1*, *CYP6Z3* and *CYP6Z4*. Unlike traditional neurotoxic insecticides, chlorfenapyr belongs to the pyrrole class and functions as a pro-insecticide that requires P450-mediated bioactivation to exert its toxic effect through the disruption of oxidative phosphorylation in the mitochondria (Black et al., 1994; Guengerich and Mitchell, 1980; Oxborough et al., 2019). Our data suggest that *CYP6Z* enzymes, in addition to their detoxifying role for pyrethroids and carbamates, may also be capable of metabolizing chlorfenapyr into its active form, or at least participating in its activation. This aligns with previous findings indicating that certain mosquito cytochrome P450 enzymes particularly anopheles *CYP6M2* and *CYP6P3* can enhance the bioactivation of chlorfenapyr (Yunta et al., 2019; Yunta et al., 2023). Similarly, previous study have shown that transgenic expressing of *An. funestus* pyrethroids resistant gene *CYP6P9a/b* enhanced flies susceptibility to chlorfenapyr (62–77% mortality) compared to the control (38%), while RNAi knockdown of this same *CYP6P9a* gene in field resistant *An. funestus* restores chlorfenapyr efficacy (Tchouakui et al., 2024).

This observation reinforces the rationale for integrating chlorfenapyr-based interventions, such as Interceptor® G2 a dual-active ITN combining chlorfenapyr and alpha-cypermethrin, into malaria vector control programs, particularly in areas where P450-mediated resistance to pyrethroids is widespread (Assenga et al., 2025) (Huang et al., 2023; WHO, 2023). In such contexts, P450 overexpression might paradoxically enhance the efficacy of chlorfenapyr through increased metabolic activation, a phenomenon previously reported in *An. gambiae* and *An. funestus* populations (Oxborough et al., 2015; Yunta et al., 2016). This finding adds a compelling dimension to the use of chlorfenapyr as a pyrethroid resistance-breaking tool, particularly in combination nets or in rotational IRS programs, where it can help mitigate the spread and impact of pyrethroid resistance (Che-Mendoza et al., 2021; Minwuyelet et al., 2025). However, further metabolic assays using recombinant *CYP6Z* proteins are needed to confirm the nature and efficiency of chlorfenapyr bioactivation by this gene family. Understanding the kinetics and structural interaction of *CYP6Z* gene family with chlorfenapyr will be essential for predicting efficacy and avoiding potential antagonistic effects when used in combination with other insecticides.

## 5. Conclusion

This study provides robust functional evidence that the *CYP6Z* gene subfamily is a central driver of broad-spectrum metabolic resistance in *Anopheles gambiae* s.s. through increased gene expression. By integrating transcriptomic, genomic, structural, and metabolic analyses with both *in vitro* and *in vivo* validation, we show that *CYP6Z* genes work in concert to metabolize a wide range of insecticide classes, including pyrethroids, organophosphates, carbamates, and neonicotinoids. At the same time, these enzymes increase susceptibility to the pro-insecticide chlorfenapyr, an important ingredient in use in the next generation LLINs. Our findings clarify the molecular mechanisms underlying cross-resistance in malaria vectors and reveal a critical vulnerability that can be strategically targeted with pro-insecticides. This work highlights the importance of ongoing surveillance of P450-based metabolic resistance and supports the rational use of new vector control tools to address the growing challenge of insecticide resistance and sustain malaria control efforts.

## Data availability

All data generated or analysed in the present study are included in the article and its Additional files. The datasets derived from the RNAseq are accessible on PRJEB97187. The DNA sequences of *CYP6Z3* reported in this paper have been deposited in the GenBank database accession numbers PX684052 - PX684093.

## Authors contributions

Conceptualization; C.S.W and S.S.I.; Study design M.F.M.K, S.S.I., and C.S.W; data curation, S.S.I., and C.S.W.; formal analysis, M.F.M.K., A.T.; A.M., and S.S.I.; funding acquisition, C.S.W and S.S.I.; investigation, M.F.M.K., A.T.; A.M., and S.S.I.; methodology, M.F.M.K.;, A.T.; V.B.N; J.A.K.; A.F.N., B.N.K., M.W.; H.I., S.S.I and C.S.W.; software, M.F.M.K., A.T.; A.M., J.A.K.; S.S.I., and C.S.W.; supervision, S.S.I., and C.S.W.; validation, S.S.I. and C.W.; writing—original draft, M.F.M.K.; writing—review and editing, M.F.M.K., A.T.; S.S.I., and C.S.W. All authors have read and agreed to the published version of the manuscript.

## Acknowledgements

We are grateful to the collection sites communities for allowing us access to their sites for mosquito’s collection. We are also grateful the following persons; Fistel J. Woume, Marceline Nke M., Army F. Ngong, Brandon N. Kongnyu and Jonathan Chedjoun for their tremendous help with bench work.

## Consent for publication

Not Applicable.

